# Layer 6 is a hub for cholinergic modulation in the mouse auditory cortex

**DOI:** 10.1101/2025.06.09.658740

**Authors:** Lucas G. Vattino, Kameron K. Clayton, Troy A. Hackett, Anne E. Takesian, Daniel B. Polley

## Abstract

Basal forebrain cholinergic neurons (BFCNs) densely innervate auditory cortex (ACtx), conveying signals linked to internal brain states and external sensory cues. Several studies have shown that acetylcholine (ACh) rapidly modifies local cortical circuits via nicotinic ACh receptors (nAChRs) on layer 1 (L1) inhibitory neurons. BFCN terminals are also abundant in L6, though the mechanisms and functional consequences of cholinergic modulation in deeper cortical layers has received less attention. Here, we performed multi-plex *in situ* labeling across cortical layers and cell types and found that L6 pyramidal neurons (L6-PNs) were highly enriched in diverse nAChR subunit and muscarinic ACh receptor (mAChR) transcripts. *In vivo* optogenetic activation of BFCN axons revealed persistent modulation of regular spiking (RS) units in L2-6 but a rapid phasic activation only in L6. In acute slices, optogenetic activation of BFCN axons elicited fast excitatory post-synaptic potentials via nAChRs in L6-PNs, comparable to responses in L1-INs, and slower inhibitory responses mediated by mAChRs. These findings identify L1 inhibitory neurons and L6 excitatory neurons as two hubs that mediate BFCN modulation of cortical circuits. Transcriptional, synaptic, and local circuit connectivity differences between L1 and L6 hubs may allow BFCN inputs to shape perception and plasticity on distinct timescales.

## Introduction

Acetylcholine (ACh) is a critical neuromodulator that enhances the detection of salient sensory cues (Parikh et al. 2007; Goard and Dan 2009; Pinto et al. 2013; Gritton et al. 2016) and promotes experience-dependent plasticity mechanisms (Kilgard and Merzenich 1998; Froemke et al. 2007, 2013; Verhoog et al. 2016; Takesian et al. 2018; Yaeger et al. 2019) that support learning and memory (Bakin and Weinberger 1996; Froemke et al. 2007; Letzkus et al. 2011; Sabec et al. 2018; Guo et al. 2019; Asokan et al. 2023). In the rodent auditory cortex (ACtx), both primary and secondary areas receive dense projections from basal forebrain cholinergic neurons (BFCNs) located in the caudal tail of substantia innominata (SI) and the globus pallidus externus with additional projections from more caudal and ventral regions of SI that is often labeled nucleus basalis (Kim et al. 2016; Chavez and Zaborszky 2017). Caudal BFCNs exhibit sound-evoked and behaviorally-gated activation patterns at multiple timescales – from fast, transient bursts to sustained activity– to shape sensory processing and plasticity in a context-dependent manner (Letzkus et al. 2011; Eggermann et al. 2014; Fu et al. 2014; Hangya et al. 2015; Nelson and Mooney 2016; Reimer et al. 2016; Kuchibhotla et al. 2017; Crouse et al. 2020; Robert et al. 2021; Zhu et al. 2023; Kimchi et al. 2024).

The impact of ACh on cortical function depends not only on the timing and pattern of ACh release but also on the distribution and subtype diversity of its receptors across the cortical layers. Ionotropic nicotinic ACh receptors (nAChRs) and G protein-coupled muscarinic ACh receptors (mAChRs) exhibit varying affinities, kinetics and downstream intracellular signaling cascades, enabling ACh to exert both phasic and tonic effects on cell excitability, functional synaptic connectivity and plasticity (Disney and Higley 2020; Sarter and Lustig 2020). While recent studies have largely focused on the influence of ACh on superficial inhibitory microcircuits, less is known about cholinergic modulation within deeper cortical layers.

Cholinergic axons densely innervate the deepest layer of cortex across both sensory and non-sensory cortices (Mechawar et al. 2000; Bloem et al. 2014; Allaway et al. 2020). In ACtx, layer 6 pyramidal neurons (L6-PNs) receive monosynaptic input from cholinergic neurons in the BFCN (Clayton et al. 2021). L6-PNs have recently been recognized as key nodes in regulating cortical output and perceptual behavior, mediating sensory gain control and shaping cortical representation of sensory stimuli (Olsen et al. 2012; Bortone et al. 2014; Crandall et al. 2015; Guo et al. 2017; Williamson and Polley 2019; Voigts et al. 2020; Clayton et al. 2021). These L6-PNs comprise distinct subpopulations based on their projection targets (Harris and Shepherd 2015). In the ACtx, intratelencephalic L6-PNs communicate with regions such as the neocortex, striatum, and amygdala (Prieto and Winer 1999; Winer and Prieto 2001), extratelencephalic projections from small PNs at the white matter border of L6 innervate the outer shell of the inferior colliculus (Schofield 2009; Yudintsev et al. 2021), and corticothalamic (CT) L6-PNs provide feedback to the ipsilateral thalamus (Prieto and Winer 1999; Guo et al. 2017; Clayton et al. 2021). CT L6-PNs might have a role in silencing thalamocortical projections (Kim et al. 2014), but despite their privileged position at the interface of sensory input and cortical output, the mechanisms by which ACh influences these L6-PNs remain unresolved.

Here, we mapped ACh receptor heterogeneity across the cortical layers and found that nAChR and mAChR transcripts are enriched within L6-PNs. Activation of BFCN axons *in vivo* induced both transient and persistent changes in the firing rate of L6 regular spiking (RS) units, in agreement with the co-expression of diverse nAChR and mAChR subtypes. Using an acute slice preparation, we found that BFCN axon stimulation elicited both nAChR-mediated depolarizing and mAChR-mediated hyperpolarizing responses in L6-PNs of ACtx slices. Together, these findings identify L6 of the mouse primary ACtx as a hub for cholinergic modulation and reveal a mechanism by which ACh may shape cortical output during auditory processing.

## RESULTS

### Diverse nicotinic and muscarinic ACh receptor subtypes are expressed in cortical L6

Cholinergic axons innervate all cortical layers of sensory cortex, with the highest densities observed in the superficial and deep layers (Mechawar et al. 2000; Allaway et al. 2020). To investigate the laminar specificity of cholinergic modulation in ACtx, we first quantified the distribution of muscarinic and nicotinic cholinergic receptor (mAChR and nAChR) transcripts across cortical layers and cell types in primary ACtx (A1). Corticothalamic (CT) projection neurons in ACtx were labeled by injecting the retrograde tracer cholera toxin subunit-B (CTB) into the medial geniculate body (MGB), and primarily concentrated in L6 (Prieto and Winer 1999; Guo et al. 2017; Williamson and Polley 2019; Clayton et al. 2021) (**Figure 1A**, **Methods**).

**Figure 1.**
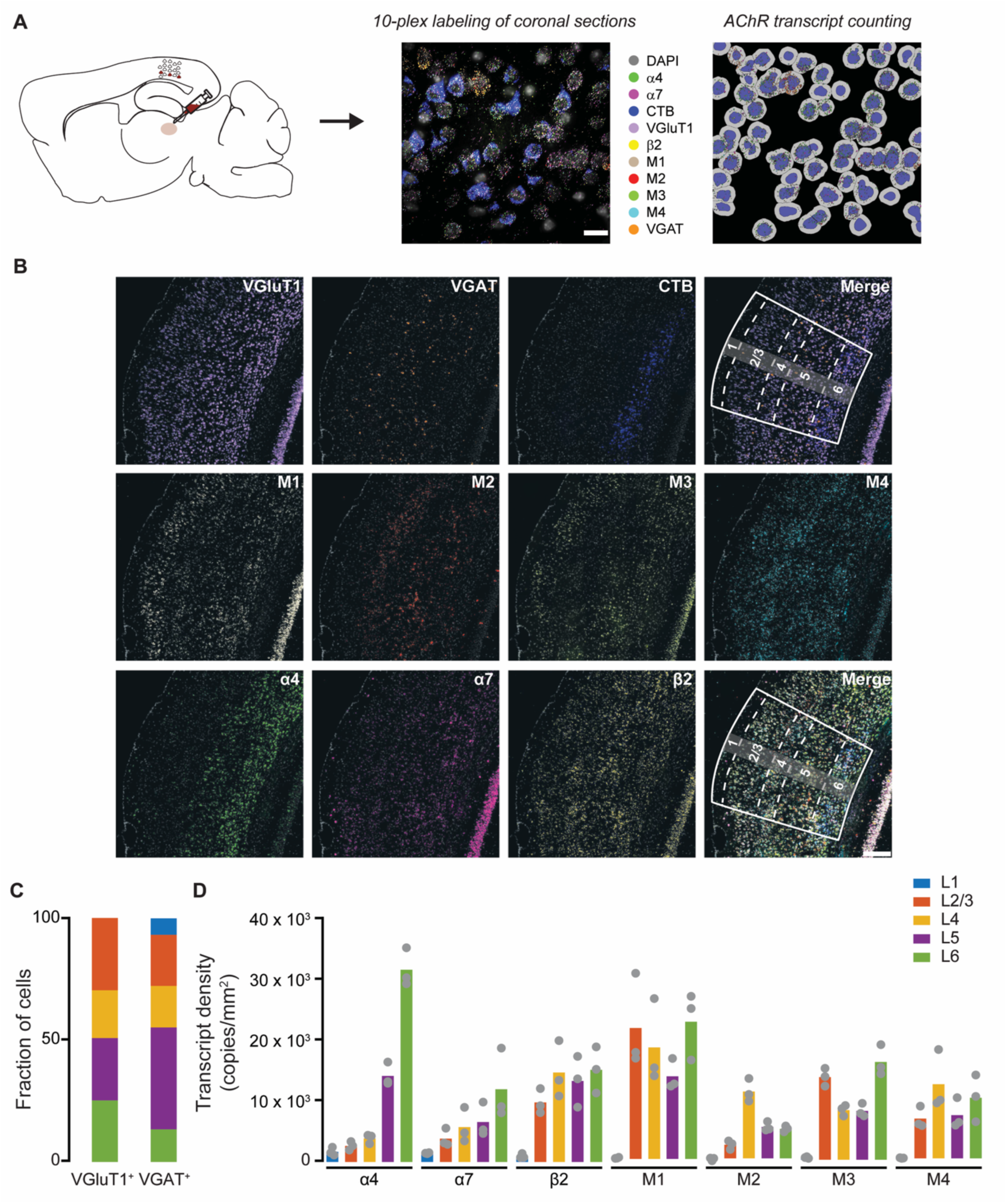
nAChR and mAChR transcripts are expressed across all layers in A1. **(A)** Schematic of the multiplex fluorescence *in situ* hybridization (FISH) strategy. **Left**: Mice (N = 3) were injected with CTB-Alexa Fluor 647 in the right MGB to label thalamic-projecting cortical pyramidal neurons (CT-PNs). **Middle**: 10-plex FISH localized mRNA transcripts encoding nAChRs and mAChRs within excitatory (vesicular glutamate transporter 1; VGluT1+) and inhibitory (vesicular GABAergic transporter; VGAT+) neurons. **Right:** Individual AChR transcripts were counted within the cytoplasm of excitatory and inhibitory neurons. Scale bar: 20 μm. **(B)** Images of A1 sections showing multiplex FISH labeling of excitatory (VGluT1^+^) and inhibitory (VGAT^+^) neurons, and mRNA transcripts encoding nAChR subunits (ɑ4, ɑ7, β2) and mAChRs (M1-4). Scale bar: 200 μm. Laminar boundaries are indicated by dashed lines. **(C)** Distribution of excitatory (VGluT1^+^, n = 2947) and inhibitory (VGAT^+^, n = 410) neurons across ACtx layers. **(D)** Quantification of the density of mRNA transcripts encoding nAChRs and mAChRs across ACtx layers.

We performed multiplex fluorescence *in situ* hybridization (FISH) to localize mRNA transcripts encoding nAChRs and mAChRs within excitatory (vesicular glutamate transporter 1; VGluT1^+^) and inhibitory (vesicular GABAergic transporter; VGAT^+^) neurons (**Figure 1A**-**C**). Transcripts for the ɑ4, ɑ7 and β2 nAChR subunits and M1-4 mAChR receptors were widely expressed across ACtx (**Figure 1B**,**D**; **Supplementary Table 1**). Notably, we observed enriched expression of both nAChRs and mAChR subunit transcripts within L6, particularly ɑ4 nAChR subunit transcripts, consistent with findings in rat ACtx (Ghimire et al. 2020).

The predominant nAChR subtypes in cortex are the homomeric α-bungarotoxin (α-Bgtx)- sensitive nAChR, composed of five α7 subunits, and the heteromeric α-Bgtx-insensitive nAChR, composed of α4 and β2 subunits (Radnikow and Feldmeyer 2018; Zoli et al. 2018). We quantified the presence of transcripts encoding for these subunits within VGAT^+^ and VGluT1^+^ populations (**Figure 2A**). Inhibitory interneurons in cortical L1 (L1-INs) are a major target for ACh across neocortical areas, mostly acting via nAChRs (Letzkus et al. 2011; Takesian et al. 2018). Consistently, we found that the majority (75.17 ± 1.19 %) of VGAT^+^ inhibitory neurons in L1 expressed mRNA transcripts encoding the ɑ7, ɑ4 and β2 subunits, likely producing homomeric ɑ7 and heteromeric ɑ4β2 nAChRs (**Figure 2B**; **Supplementary Table 2**). Across other layers, no more than 25% of VGAT^+^ neurons expressed transcripts encoding putative ɑ7 and ɑ4β2 nAChRs (**Figure 2B**; **Supplementary Table 2**). Among the VGluT1^+^ populations, those in L5 and L6 showed the highest fraction of neurons expressing transcripts encoding both ɑ7 and ɑ4β2 nAChRs or only the ɑ4β2 nAChR (**Figure 2B**; **Supplementary Table 2**). Within the CTB^+^ population of CT L6-PNs, the ratio of neurons expressing these nAChR subunit transcripts was modestly elevated as compared to the CTB^-^ population (**Figure 2C**; **Supplementary Table 3**). Similarly to what has been observed in the rat ACtx (Ghimire et al. 2020), we found a subpopulation of VGAT^+^ cells spanning L2-5 that might express ɑ7β2 nAChRs.

**Figure 2.**
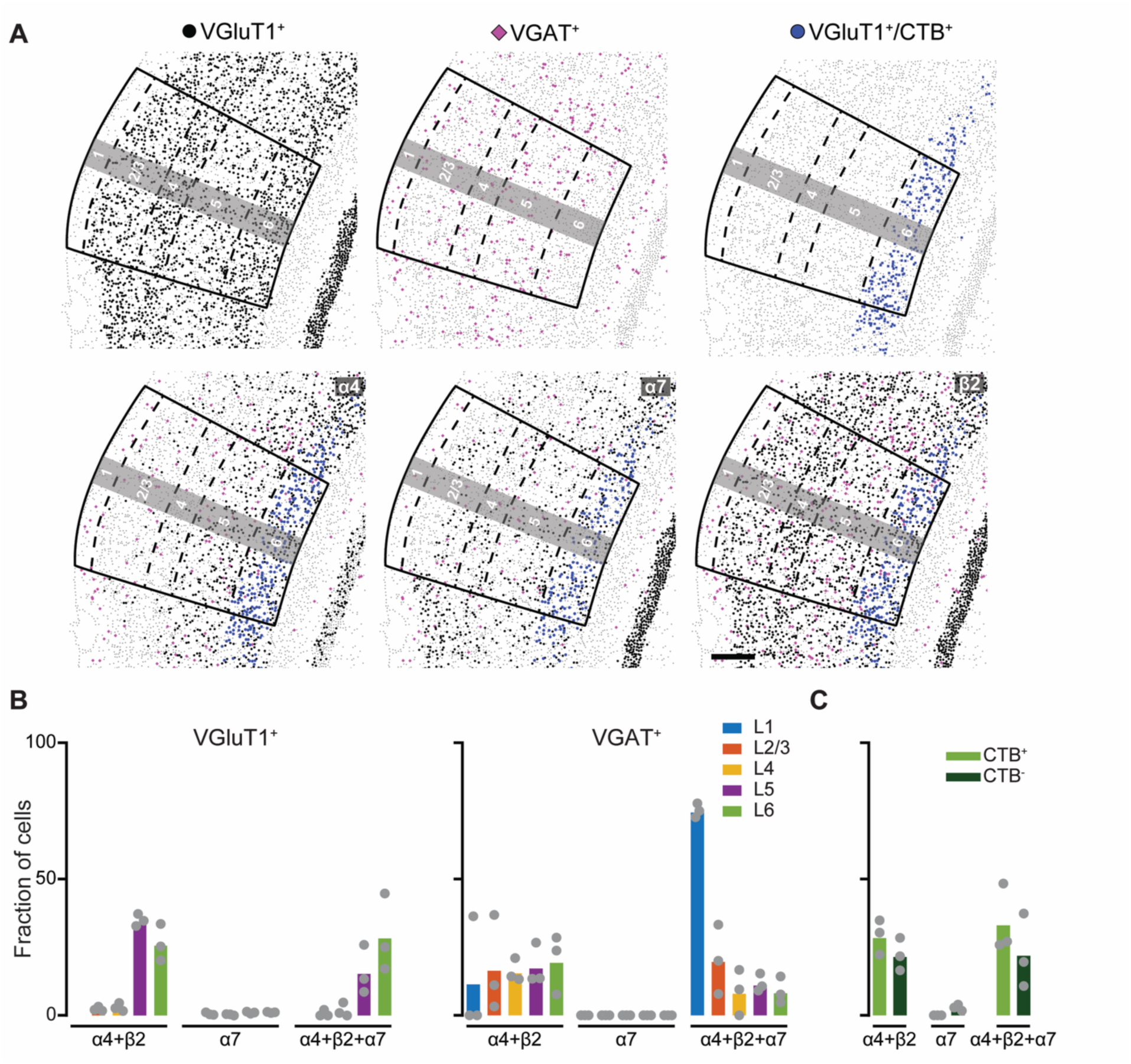
nAChR subunit transcripts are enriched in L6 of primary A1. (A) **Plots of cells** containing transcripts for each nAChR subunit combination in VGluT1^+^, VGAT^+^ and VGluT1^+^/CTB^+^ cells. Laminar boundaries are indicated by dashed lines. Scale bar: 200 μm. **(B)** Fraction of excitatory (VGluT1^+^, left) and inhibitory (VGAT^+^, right) neurons expressing subunit transcript combinations to putatively form heteromeric ɑ4β2 and homomeric ɑ7 nAChRs across ACtx layers. **(C)** Fraction of CT L6-PNs (CTB^+^) or non-CT L6-PNs (CTB^-^) expressing nAChR subunit transcripts.

We next analyzed the expression of mAChRs M1-4 transcripts. A higher proportion of VGAT^+^ inhibitory neurons in L6 expressed M2, M3 and M4 receptor transcripts as compared to inhibitory neurons within the other cortical layers. The majority of VGluT1^+^ neurons across all layers of ACtx expressed mRNAs for the M1 receptor (**Figure 3A**,**B**; **Supplementary Table 2**). We found that more neurons within the CTB^+^ population of CT L6-CNs expressed M1, M3 and M4 receptor transcripts, whereas more CTB^-^ neurons expressed M2 (**Figure 3C**; **Supplementary Table 3**). Together, these transcriptomic data reveal that both excitatory and inhibitory neurons in L6 express diverse nAChR and mAChR subtypes, highlighting L6 as a key target for ACh modulation.

**Figure 3.**
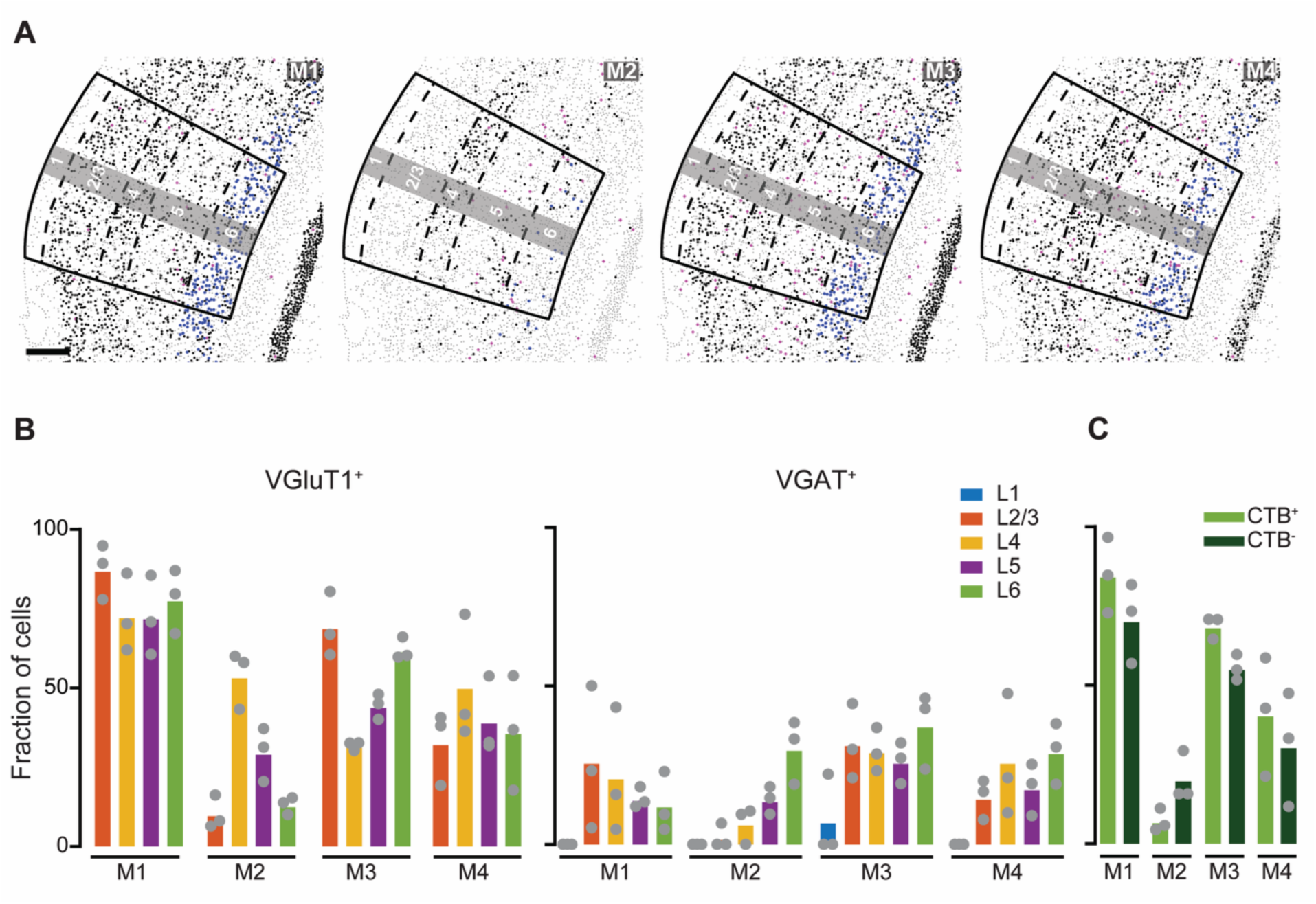
Robust expression of mAChR subunit transcripts across A1. **(A)** Plots of VGluT1^+^, VGAT^+^ and VGluT1^+^/CTB^+^ cells containing mAChR transcripts. Laminar boundaries are indicated by dashed lines. Scale bar: 200 μm. **(B)** Fraction of excitatory (VGluT1^+^, left) and inhibitory (VGAT^+^, right) neurons expressing the mAChR subtype transcripts (M1-4) across ACtx layers. **(C)** Fraction of CT L6-PNs (CTB^+^) or non-CT L6-PNs (CTB^-^) expressing mAChR transcripts.

### Cholinergic input modulates the activity of regular-spiking (RS) neurons in cortical L6

To probe the functional impact of cholinergic inputs in ACtx neurons across the cortical layers, we selectively expressed channelrhodopsin-2 (ChR2) in basal forebrain cholinergic neurons (BFNCs) using AAV2/5-Ef1a-ChR2-EYFP injections into ChAT-IRES-Cre(Δneo) x Cdh23 mice (**Figure 4A**), which offer excellent hearing into adulthood and selective expression of Cre-recombinase in cholinergic neurons (Robert et al. 2021). Four weeks post injection, we obtained translaminar recordings from primary ACtx of awake, head-fixed mice while optogenetically stimulating BFCN axons (**Figure 4A**, **B**).

**Figure 4.**
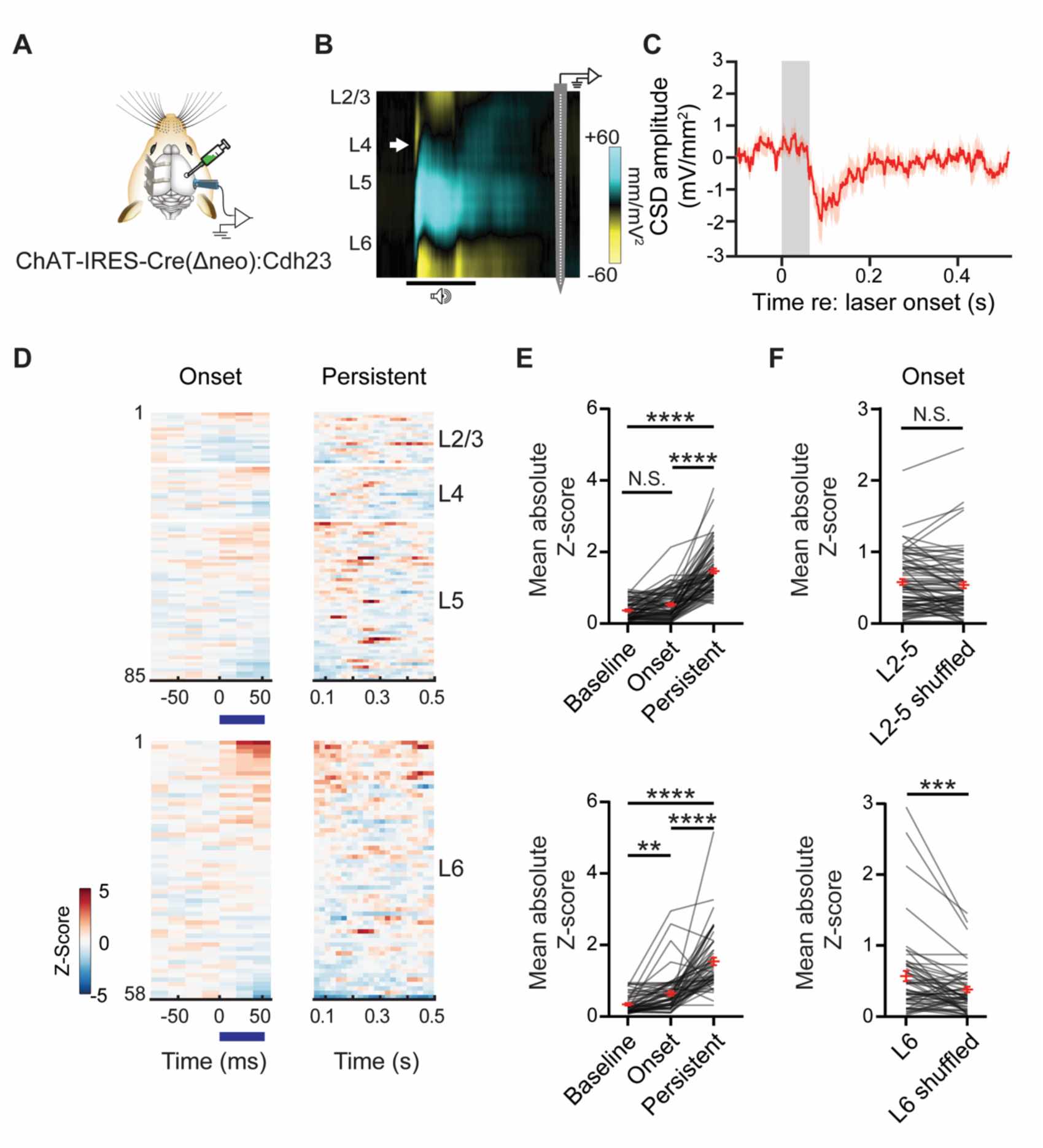
BFCN inputs modulate ACtx L6 excitatory neurons *in vivo*. **(A)** Schematic of the experimental strategy for *in vivo* single-unit recordings. Cre-dependent ChR2 was injected in the BFCN of ChAT-IRES-Cre(Δneo) x Cdh23 mice (N = 5 mice). Neural activity was recorded with a 64-channel linear silicon probe. BFCN axon terminals were stimulated with an optic fiber coupled to a 473 nm diode laser. **(B)** Extracellular recordings were made across L2/3- 6 in ACtx. The current source density (CSD) of neural activity in response to a 50 ms noiseburst was used to approximate depth for each recorded unit. White arrow indicates the early current sink in L4 elicited by the stimulus. **(C)** A brief laser pulse (50 ms, 30 mW) was used to determine the presence of the L2/3 sink in the CSD evoked by cholinergic activation (Guo et al. 2019). **(D)** Neurograms representing the Z-scored responses of RS units to a laser pulse (50 ms, 30 mW) used to activate the BFCN axons in ACtx. For each layer, units are sorted by mean Z-score responses during the laser stimulus. **(E)** Mean absolute Z-score values for units in L2-5 (**top**) or in L6 (**bottom**) calculated during Baseline (L2-5: 0.37 ± 0.03; L6: 0.34 ± 0.03), the 50 ms the laser stimulation lasted (Onset; L2-5: 0.53 ± 0.04; L6: 0.65 ± 0.03) or during the following 500 ms (Persistent; L2-5: 1.47 ± 0.07; L6: 1.54 ± 0.11). Friedman test with Dunn’s post-hoc comparison for L2-5 (**top**): Baseline vs Onset (p = 0.07); Baseline vs Persistent (p < 0.0001); Onset vs Persistent (p < 0.0001). Friedman test with Dunn’s post-hoc comparison for L6 (**bottom**): Baseline vs Onset (0.0011); Baseline vs Persistent (p < 0.0001); Onset vs Persistent (p < 0.0001). (**F**) Mean absolute Z-score values obtained during the Onset time period and compared to a shuffled control population for L2-5 (**top**; 0.55 ± 0.04 and 0.51 ± 0.05) and L6 (**bottom**; 0.54 ± 0.07 and 0.34 ± 0.04). Wilcoxon test for L2-5 vs L2-5 shuffled (p = 0.13) and L6 vs L6 shuffled (p = 0.0001).

Cortical layer boundaries were identified using current source density (CSD) profiles evoked by a 50 ms broadband noise burst, with a characteristic sink demarcating the boundary between L4 and L5 (**Figure 4B**, **Methods**). BFCN axon activation elicited CSD responses in L2/3, confirming that activation of cholinergic axons elicited local network activity in ACtx (**Figure 4C**) (Guo et al. 2019). However, cholinergic axons innervate all layers of the cortex (Mechawar et al. 2000; Bloem et al. 2014; Allaway et al. 2020), imposing a challenge to isolate the pre- and post- synaptic contributors of the BFCN-evoked CSD.

Although cholinergic inputs to the neocortex are generally viewed as modulatory and therefore unlikely to directly elicit spiking, the abundance of nAChRs in deep layer PNs suggested that phasic ACh release could play a “driver” role by evoking spikes rather than just modulating local network activity. To address this possibility, we made extracellular recordings with translaminar probes and isolated single units in L2-6 with regular spiking (RS) waveforms that likely identify excitatory PNs (**Methods**). RS units exhibited heterogeneous responses to BFCN axon stimulation, with some units showing an increase in spiking activity and others a suppression of activity (**Figure 4D**). We quantified these effects during laser stimulation (onset) and after the laser was turned off (persistent) by comparing the absolute mean z-scored activity of each unit during both time periods (**Figure 4D**, **E**; **Methods**). While all layers displayed delayed, persistent BFCN-driven modulation 100-500 ms after laser stimulation, we observed that some L6 RS units showed time-locked spiking activity during the 50 ms laser stimulation window (**Figure 4D**, **E**). To further confirm these findings, we compared the onset absolute mean z-scored activity of each unit to a control obtained by randomly shuffling the time bins of the unit’s neurogram (**Figure 4F**; **Methods**) and found that stimulation of BCFN axons elicited significant changes in L6 RS spiking, but not in other layers. These findings demonstrate that BFCN inputs modulate L6 RS units in primary ACtx and may act through both fast and slow receptor mechanisms in a layer-specific manner.

### L6-PNs exhibit mAChR- and nAChR-mediated responses in vitro

The development of tools to study BFCN signaling in the neocortex have made it possible to better understand the *in vivo* function of cholinergic inputs, but the limited spatiotemporal resolution precludes any insight into the layer-specific effects and pre- versus post-synaptic mechanisms of ACh action (Ananth et al. 2023). To resolve the receptor-specific contributions underlying fast and sustained cholinergic effects in L6 and to confirm that BFCNs have monosynaptic effects on L6 PNS, we turned to an *in vitro* approach. We performed whole-cell patch-clamp recordings from L6-PNs in acute primary ACtx slices during optogenetic stimulation of BFCN axons (**Figure 5A**). Glutamatergic AMPA and NMDA receptors were blocked to isolate cholinergic effects.

**Figure 5.**
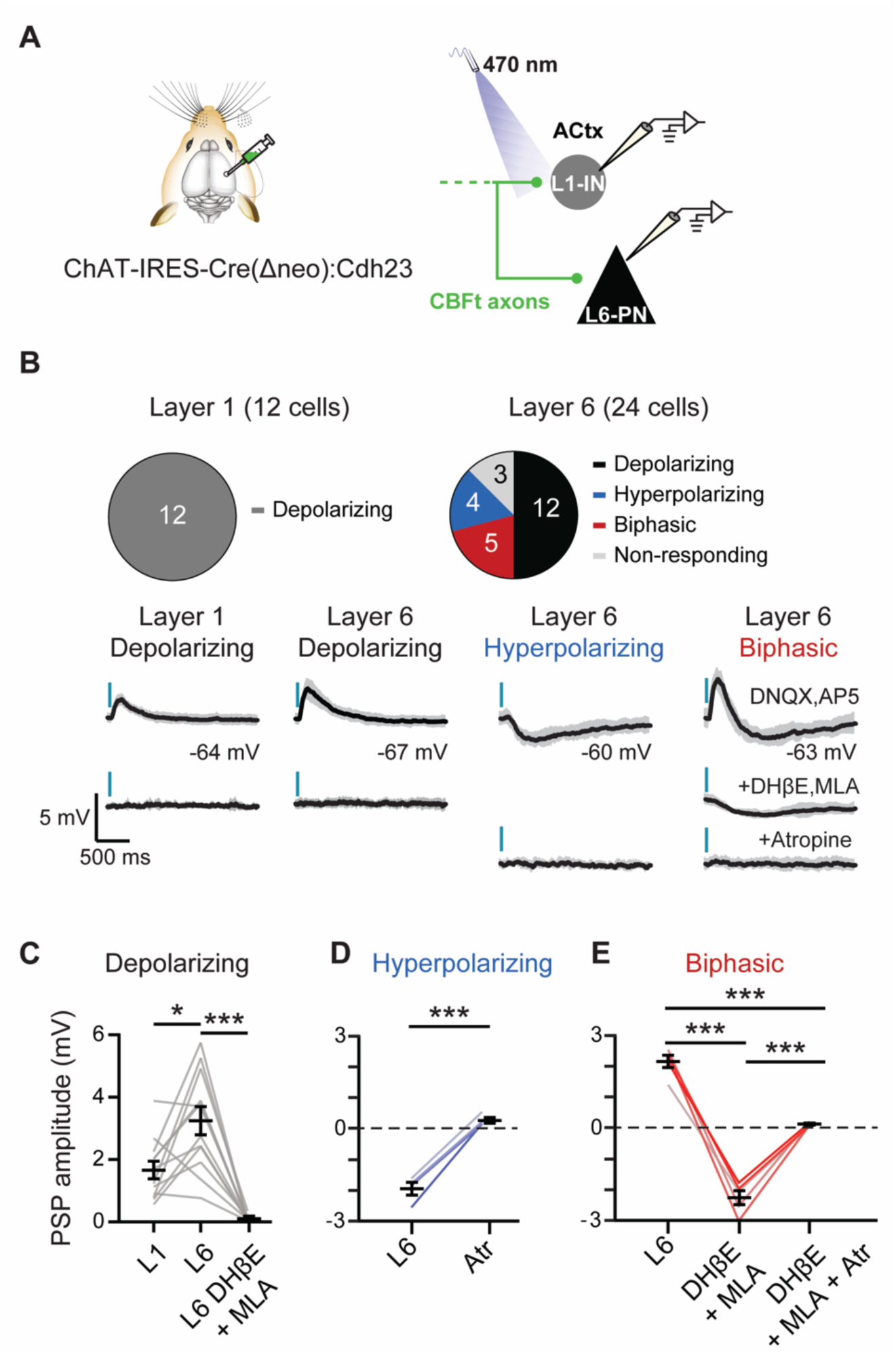
BFCN axon stimulation evokes mAChR- and nAChR-mediated postsynaptic potentials in L6-PNs neurons. **(A)** Schematic of the experimental strategy. Cre-dependent ChR2 was injected into the BFCN of ChAT-IRES-Cre(Δneo) x Cdh23 mice (N = 6 mice). BFCN axons in ACtx were stimulated with a 470 nm LED (14 mW/mm^2^, 5 ms) and whole-cell current clamp recordings were obtained from L1-INs and L6-PNs. **(B)** Postsynaptic potentials (PSPs, mean ± SD of 10 trials) recorded in the presence of the AMPA and NMDA receptor blockers (DNQX; 20 μM, AP5; 50 μM) revealed heterogeneous responses in L6-PNs to activation of BFCN axons. **(C)** The amplitudes of depolarizing PSPs in L6-PNs (**middle**; 3.25 ± 0.45 mV) were significantly larger than those in L1-INs within the same slices (**left**; 1.66 ± 0.28 mV). These responses were eliminated by nAChR antagonists DHβE (10 μM) and MLA (10 μM) (**right**; 0.09 ± 0.03 mV). Wilcoxon test L1 vs L6: p = 0.012; Wilcoxon test L6 vs L6 DHβE + MLA: p = 0.0005. **(D)** L6-PNs showing hyperpolarizing PSPs (-2.05 ± 0.21 mV) eliminated by the mAChR antagonist atropine (Atr, 10 μM; 0.18 ± 0.09 mV). Paired t-test L6 vs Atr: p = 0.001. **(E)** Biphasic PSPs in L6- PNs (2.15 ± 0.20 mV) eliminated by sequential nAChR blockade using DHβE (10 μM) and MLA (10 μM) (**middle**; -2.27 ± 0.23 mV) and mAChR blockade using atropine (Atr, 10 μM; **right**; 0.12 ± 0.03 mV). RM-ANOVA with post-hoc Fisher’s comparison: L6 vs L6 DHβE + MLA (p = 0.0002); L6 vs L6 DHβE + MLA + Atr (p = 0.0004); L6 DHβE + MLA vs L6 DHβE + MLA + Atr (p = 0.0005).

BFCN activation evoked heterogeneous postsynaptic responses in L6-PNs (**Figure 5B**). Among the recorded neurons, 50% exhibited depolarizing postsynaptic potentials (PSPs) that were eliminated by nAChR antagonists MLA and DHβE (**Figure 5B**, **C**). These depolarizing responses were significantly larger than those observed in L1-INs recorded within the same slices (**Figure 5B**, **C**). A subset (20%) of L6-PNs showed exclusively hyperpolarizing PSPs that were abolished by the mAChR antagonist atropine (**Figure 5B**, **D**). Additionally, 16% of L6-PNs exhibited biphasic responses consisting of an initial depolarizing PSP followed by a sustained hyperpolarization, which were sequentially abolished by nAChR and mAChR blockade, respectively (**Figure 5B**, **D**).

Together, our findings indicate that L6-PNs are key nodes of direct cholinergic input mediated both by fast-acting ionotropic nAChRs and slower-acting metabotropic mAChRs. The combination of depolarizing and hyperpolarizing responses indicates that cholinergic inputs exert temporally precise and functionally complex effects on L6 circuitry within the ACtx.

## DISCUSSION

Decades of research have established acetylcholine as a key modulator of cortical processing and a driver of plasticity, yet the precise cellular and circuit mechanisms through which it operates remain incompletely understood (Picciotto et al. 2012; Obermayer et al. 2017). Our findings identify cortical layer 6 (L6) as a major hub for cholinergic modulation, where pyramidal neurons exhibit a diverse, functionally complex repertoire of AChR signaling. Although prior studies have largely focused on superficial cortical layers (Letzkus et al. 2011; Arroyo et al. 2012; Poorthuis et al. 2018; Takesian et al. 2018), our data show that L6-PNs, including corticothalamic (CT) neurons, are robustly modulated by cholinergic input from the BFCN.

Neocortical cholinergic signaling has been traditionally characterized as slow and volume-mediated, although a revised model has emerged in which both fast and slow ACh-mediated transmission co-exist to shape cortical states and behavior across timescales (Sarter et al. 2009; Disney and Higley 2020; Sarter and Lustig 2020). The spatiotemporal effects of cholinergic inputs are determined by the receptor subtype involved (Higley and Picciotto 2014), with ionotropic nAChRs mediating fast, synaptic-like excitatory transmission (Albuquerque et al. 2009) and G-protein coupled (GPCR) mAChRs inducing delayed and prolonged excitatory and inhibitory effects (Thiele 2013). Using transcriptomic profiling and functional recordings, we show that L6- PNs express a broad array of both nAChRs and mAChRs. Notably, L6-PNs are enriched for RNA transcripts of ɑ7, ɑ4 and β2 subunits, likely producing ɑ7 and ɑ4β2 nAChRs – the two most prevalent nAChRs in cortex (Zoli et al. 2018; Ghimire et al. 2020). Indeed, ∼50% of the L6-PNs express transcripts that encode for subunits that can combine and form homomeric ɑ7 and heteromeric ɑ4β2 nAChRs, a higher fraction than any other cortical excitatory population in our study (**Figure 2**). These L6-PNs also robustly express transcripts encoding M1-M4 mAChRs (**Figure 3**) that primarily signal via Gq/11 (M1,3) or Gi/o (M2,4) protein pathways (Thiele 2013). The downstream effects of these GPCRs vary by cell type and depend on proximity to signaling molecules and ion channels, resulting in either depolarization or hyperpolarization (Thiele 2013). A striking example is the M1 receptor, which can induce either closure of K^+^ channels to excite neurons (Womble and Moises 1992; Ghamari-Langroudi and Bourque 2004; Carr and Surmeier 2007; Giessel and Sabatini 2010) or opening of SK Ca^2+^-activated K^+^ channels to inhibit neurons (Brombas et al. 2014). Thus, receptor expression alone cannot predict postsynaptic effects of cholinergic modulation.

The cholinergic receptor diversity in L6-PNs underlies the heterogenous responses to BFCN axon stimulation observed *in vitro* (**Figure 5**) – ranging from fast, phasic depolarization via nAChRs to delayed hyperpolarization via mAChRs. These dual responses were even observed within single neurons, highlighting a convergence of receptor signaling that may enable flexible, time-dependent modulation of excitability. Consistently, *in vivo* single unit recordings revealed that BFCN axon activation can bidirectionally modulate RS L6 units in ACtx on distinct timescales, reflecting the co-engagement of fast and slow cholinergic mechanisms (**Figure 4**). Some RS L6 units showed phasic spiking, suggesting that ACh can act as a rapid, synaptic-like ‘driver’ through nAChRs – paralleling mechanisms observed in L1 interneurons. In contrast, more sustained cholinergic effects observed across layers, including L6, may be mediated by mAChRs. These results support a growing model that cortical ACh signaling in L6 operates through temporally and functionally distinct modes that interact to support dynamic cortical states.

The diverse complement of cholinergic receptors expressed in L6-PNs may enable activity-dependent modulation by ACh. For example, ACh transiently excites neurogliaform neurons in superficial cortex via nAChRs at rest, but inhibits these neurons for long time periods via mAChRs during active firing states (Brombas et al. 2014). Analogously, L6-PNs may show distinct cholinergic responses depending on their membrane potential or recent activity history, positioning ACh as a context-dependent modulator of deep-layer excitability.

The modulatory effects of cholinergic axons are also shaped by an interaction between presynaptic firing patterns and the composition of postsynaptic ACh receptors (Laszlovszky et al. 2020; Schlingloff et al. 2025). Cholinergic neurons exhibit a range of firing dynamics -- from spikes triggered by aversive events such as air puffs (Hangya et al. 2015) to responses to sound stimuli that rapidly scale in amplitude as animals learn that they predict environmental threats (Kuchibhotla et al. 2017; Guo et al. 2019; Crouse et al. 2020; Robert et al. 2021). These firing patterns could influence the accumulation of ACh, potentially determining which receptor subtypes are activated. High-frequency firing may produce more sustained, elevated ACh levels that could recruit lower-affinity extrasynaptic receptors and mediate modulatory effects over extended timescales. Conversely, lower ACh concentrations could preferentially activate high-affinity, synaptic nAChRs and mAChRs (Hay et al. 2016; Yang et al. 2020) mediating phasic responses in postsynaptic neurons. In the somatosensory cortex, for example, low concentrations of ACh modulated all recorded L6 pyramidal cells via mAChRs, whereas high concentrations selectivity activated CT L6-PNs via ɑ4β2 nAChRs (Qi et al. 2025). Thus, the temporal dynamics of cholinergic input may selectively engage specific L6-PNs via ACh receptor subtypes, shaping cortical output by biasing activity toward functionally distinct circuit pathways.

In the ACtx, cholinergic signaling is tightly linked to locomotion, reinforcement learning and the detection of behaviorally relevant auditory cues (Nelson and Mooney 2016; Guo et al. 2019; Robert et al. 2021). Accurate sensory detection is highly dependent on cortical state (McGinley et al. 2015), which exists along a continuum. Intermediate levels of arousal – that directly correlate with cholinergic activity in the cortex (Reimer et al. 2016) – are optimal for sensory signal detection (McGinley et al. 2015). Given their enrichment of both nAChRs and mAChRs (**Figure 2** and **Figure 3**), L6-CT neurons are well positioned to transmit ACh signals to regulate thalamic feedback. Consistent with our transcriptomic profiling, L6-CT neurons in the prefrontal cortex (PFCtx), visual and somatosensory cortices may be preferentially depolarized by ACh via the combined action of nAChRs and mAChRs, while neighboring corticocortical neurons are less responsive (Kassam et al. 2008; Sundberg et al. 2018; Yang et al. 2020; Qi et al. 2025). Moreover, a recent study in ACtx showed that L6-CT neurons are selectively modulated during active behavior by motor-related inputs that likely arise from the BFCN (Clayton et al. 2021). L6-CT neurons in sensory cortices project back to the thalamus (Prieto and Winer 1999; Harris and Shepherd 2015), providing a feedback mechanism that enables dynamic regulation of cortical gain in response to behaviorally relevant stimuli (Crandall et al. 2015; Guo et al. 2017; Clayton et al. 2021). Importantly, the precise timing of ACtx L6-CT neuron spiking relative to a sound stimulus is critical for determining whether their activity enhances or reduced the gain in the thalamus. Therefore, these neurons can act as a switch between sensory processing modes that optimize either the enhanced detection of faint stimuli or improved discrimination between similar stimuli (Guo et al. 2017). Thus, cholinergic modulation of cortical L6-CT neurons may filter thalamic inputs according to behavioral demands, supporting flexible sensory processing.

Cholinergic inputs to the ACtx not only modulate moment-to-moment encoding of the sensory environment but shape long-term plasticity (Kilgard and Merzenich 1998; Froemke et al. 2007, 2013; Takesian et al. 2018; Guo et al. 2019). These effects are highly layer-specific and depend on factors such as receptor subtype, subcellular localization, and postsynaptic neuron identity (Couey et al. 2007; Poorthuis et al. 2013; Verhoog et al. 2016). In L6 of the prefrontal cortex, activation of β2-containing nAChRs on L6-PNs enhances synaptic plasticity (Verhoog et al. 2016). In the SCtx, cholinergic signaling modulates dendritic integration in L5-PNs, giving rise to dendritic plateau potentials that can drive robust neuronal output (Williams and Fletcher 2019). Additionally, cholinergic modulation of diverse IN subtypes across layers can influence PN activity through various inhibitory circuit motifs, including feed-forward inhibition, lateral inhibition and disinhibition (Tremblay et al. 2016). Future studies should investigate the mechanisms by which ACh regulates long-term plasticity in deep-layer circuits in ACtx, particularly under behaviorally relevant conditions.

Our findings show that cholinergic signaling in the neocortex is highly circuit-specific, operating through mechanisms shaped by the distribution of receptor subtypes across cell types and cortical layers (Obermayer et al. 2017). This precision reflects the functional topography of the cholinergic basal forebrain (CBF), where anterior regions such as the horizontal limb of the diagonal band (HDB) are preferentially engaged by behavioral outcomes, whereas caudal regions within the BFCN including the globus pallidus and substantia innominata (GP/SI) preferentially respond to salient sensory stimuli and aversive events (Robert et al. 2021). Projections from these nuclei exhibit laminar specificity: in SCtx, HDB and rostral SI axons target superficial layers (e.g., L1), whereas rostral SI and NB axons preferentially innervate deeper layers (Allaway et al. 2020). A similar laminar specificity has been observed in the PFCtx (Schmitzer-Torbert et al. 2005; Clayton et al. 2021), suggesting parallel cholinergic pathways that modulate distinct microcircuits. This spatially-specific organization of cholinergic modulation across the cortical lamina supports a model in which ACh release is not uniform but may dynamically modulate separate cortical circuits in a task-dependent manner. Notably, L6-CT neurons robustly recruit PV interneurons (West et al. 2006; Frandolig et al. 2019), while L1-INs inhibit PV cells (Letzkus et al. 2011; Pi et al. 2013; Abs et al. 2018; Takesian et al. 2018; Cohen-Kashi Malina et al. 2021; Hartung et al. 2024), revealing a complex, multilayered regulatory mechanism. A subpopulation of cortical L6-PNs may also send projections to L1 interneurons (L1-INs), suggesting interlaminar communication (Ledderose et al. 2023). Together, these observations support a model in which cholinergic inputs to L1 and L6 engage distinct, and potentially competing, microcircuits that modulate sensory processing across behavioral states and timescales.

Together, our findings identify L6 of ACtx as a critical site for cholinergic modulation, where pyramidal neurons integrate fast and slow ACh signals via nAChRs and mAChRs. These interactions operate across distinct temporal and spatial scales, supporting a model in which acetylcholine dynamically sculpts cortical computations through finely tuned layer- and cell-type– specific mechanisms. This highlights the importance of incorporating deep-layer circuits into frameworks of neuromodulation and sensory plasticity.

## Acknowledgments

This work was supported by funds from the NIH NIDCD R01DC018353 (A.E.T.), R01DC017078 (D.B.P.), R01DC015388 (T.A.H.), the Nancy Lurie Mark Family Foundation (D.B.P. and A.E.T.) and the Centurion Foundation (D.B.P. and A.E.T.).

## Author contributions

D.B.P. and A.E.T. conceived this study. K.K.C. performed surgeries. T.A.H. performed and quantified histology experiments. K.K.C. collected *in vivo* electrophysiology data. K.K.C. and L.G.V. analyzed *in vivo* electrophysiology data. L.G.V. collected and analyzed *in vitro* electrophysiology data. L.G.V. and A.E.T. wrote the manuscript with input from all authors.

## METHODS

### Experimental model

All experiments were carried out in mice obtained by crossing ChAT-IRES-Cre(Δneo) males (RRID: IMSR_JAX:031661) with Cdh23 females (RRID: IMSR_JAX:018399), purchased from Jackson Laboratories. Mice were group housed and maintained under a 12:12 hr light:dark cycle, with access to food and water *ad libitum*. Both male and female adult animals (P60-90) were used for this study. All procedures were approved by the Institutional Animal Care and Use Committee at Massachusetts Eye and Ear.

### Surgeries

Mice were anesthetized with 5% isoflurane in O2 and moved to a stereotactic surgery rig where the mouse was maintained on 2% isoflurane during the procedure. A homeothermic blanket system was used to maintain body temperature at 36.5° C (FHC). After shaving and disinfecting the skin, the dorsal surface of the scalp was retracted, and the periosteum was removed. For multiplex fluorescence *in situ* hybridization (FISH) experiments, a burr hole was made in the skull (coordinates: A-P: 2.4; M-L: 2.2, depth: 2.95 mm, 3.15mm (2 depths)) to target the right medial geniculate body (MGB). A motorized injection system (Nanoject) was used to inject retrograde tracer Cholera Toxin subunit-B (CTB) Alexa Fluor-647 (Thermofisher, C34778) 2.95 and 3.15 mm below the pial surface (75 nL at each depth, 9 nL/min). For *in vitro* and *in vivo* electrophysiology experiments, a burr hole was drilled in the skull at 0.8 mm posterior from bregma and 2.7 lateral from the midline, to target the right caudal tail of the cholinergic basal forebrain (BFCN). At this location, mice were injected with 500 nL of AAV2/5-Ef1a-ChR2-EYFP (Mass Eye and Ear Viral Vector core, addgene plasmid# 35509), 3.45 mm below the pial surface at 9 nl/min. In mice used for *in vivo* electrophysiology and BFCN optogenetic stimulation, the skull surface was prepped with etchant (C&B metabond) and 70% ethanol before affixing a titanium head plate (iMaterialise) to the dorsal surface with dental cement (C&B metabond), and a ground wire (AgCl) was implanted over the left occipital cortex. All electrophysiology experiments were performed four weeks after injections. At the beginning of all recovery procedures, Buprenex (1 mg/kg) and Meloxicam (5 mg/kg) were administered and following procedures, the animal was transferred to a warmed recovery chamber.

### Histology and *in situ* hybridization

Five days after the CTB-Alexa Fluor 647 injections, mice were anesthetized with isoflurane 5% in O2, and perfused transcardially with 4% phosphate buffered paraformaldehyde (PFA). The brains were removed and postfixed for 24 hours, then transferred to 30% sucrose solution at 4° C for cryoprotection. Coronal sections (10 µm) were obtained from the rostral-caudal extent of A1 of fresh frozen brains. Multiplexed fluorescence *in situ* hybridization (FISH) was used to detect expression of nAChR subunit transcripts (β2, α4, and α7) and mAChR (M1, M2, M3, M4) transcripts in fresh frozen tissue sections (10µm) from A1 (**Figure 1**; **Table 1**). Neuronal subtypes were identified by expression of transcripts for the vesicular glutamate transporter 1 (VGluT1, *Slc17a7*) or the vesicular GABA transporter (VGAT, *Slc32a1*). Assays utilized RNAscope riboprobes, reagents, and protocols produced by Advanced Cell Diagnostics (ACD) (Hayward, CA) for multiplexed FISH, following ACD protocols and procedures previously described (Ghimire et al. 2020, 2023). Briefly, sections were post-fixed for 60 min in 4% PFA, dehydrated in an ascending ethanol series, immersed in ACD Target Retrieval solution for 5 min at 95° C, incubated for 30 min in Protease 3 solution at 40° C, followed by riboprobe hybridization (cocktail of all riboprobes) for 2 hours at 40° C. Sequential amplification steps culminated in binding of fluorescent conjugates (T1-Alexa 488, T2-Atto 555, T4-Alexa 750) to probe channels T1, T2 and T4 and counterstaining with DAPI. Sections were imaged, fluorescent tags stripped, then amplification proceeded for probes T5, T6, and T8, followed by imaging for T5 – T8, stripping, amplification for probes T9, T10, and T12, then imaging for probes T9 – T12. The T3, T7, and T11 channels were not hybridized with any probe, as that channel contained the CTB Alexa 647 signal, serving as a marker of cells labeled by retrograde CTB transport.

**Table 1.** RNAscope reagents and riboprobes for nAChRs, mAChRs and neuronal class detection.

| Cell type | Target | Gene | Cat. No. | Accession No. | Position |
| --- | --- | --- | --- | --- | --- |
| All | Nicotinic receptor subunit, $\alpha 4$ | *Chrna4* | 429871-T1 | NM_015730.5 | 1129 - 2273 |
| All | Nicotinic receptor subunit, $\alpha 7$ | *Chrna7* | 465161-T2 | NM_007390.3 | 175 - 1122 |
| Glutamatergic neurons | Vesicular glutamate transporter 1 | *Slc17a7* (VGluT1) | 416631-T4 | NM_182993.2 | 464 - 1415 |
| All | Nicotinic receptor subunit, $\beta 2$ | *Chrnb2* | 449231-T5 | NM_009602.4 | 232 - 1805 |
| All | Muscarinic receptor 1 | *Chrm1* | 495291-T6 | NM_001112697.1 | 851 - 1994 |
| All | Muscarinic receptor 2 | *Chrm2* | 495311-T8 | NM_203491.3 | 940 - 1960 |
| All | Muscarinic receptor 3 | *Chrm3* | 437701-T9 | NM_033269.4 | 353 - 1395 |
| All | Muscarinic receptor 4 | *Chrm4* | 410581-T10 | NM_007699.2 | 400 - 1330 |
| GABAergic neurons | Vesicular GABA transporter | *Slc32a1* (VGAT) | 319191-T12 | NM_009508.2 | 894 - 2037 |
| ---- | RNAscope<br>HiPlex12<br>Reagents Kit | ---- | 324409 | ---- | ---- |

### Multi-fluorescence imaging and cellular phenotyping

Images of multiplex FISH-reacted sections were obtained using a 20x objective with a Nikon 90i epifluorescence microscope and Hamamatsu Orca 4.0 CCD camera, controlled by Nikon Elements AR software. Each of the three images sets (T1 – T4, T5 – T8, T9 – T12) were imaged in 5 color channels, aligned by DAPI staining, then merged to form a resultant multiplexed image containing the three nAChR and four mAChR channels, plus CTB and DAPI (**Figure 1**). Images were imported in HALO pathology software (Indica labs, Albuquerque, NM) for analysis.

Transcript density was obtained by counting individual transcripts for each nAChR and mAChR target, quantified by cortical layer (**Figure 1**). Cells expressing the transcripts of each nAChR and mAChR target were tallied by cortical layer and neuronal class, based on co-expression with VGluT1 (glutamatergic), VGAT (GABAergic) and CTB. Cells that contained 5 or more transcripts met the threshold for tagging as positively labeled for a given probe target. Cellular phenotypes, identified based on co-expression patterns of the nAChR and mAChR targets, tallied by neuronal subtype and cortical layer (**Figure 2** and **Figure 3**).

### Single unit recordings during optogenetic stimulation in head-fixed mice

Animals were briefly anesthetized with isoflurane (5% in O2 for induction, and 2% during procedure) while a small 1 mm x 1 mm craniotomy was made along the caudal end of the right temporal ridge to expose A1 (1.5 mm rostral from lambda). A small chamber was built around the craniotomy with UV-cured cement and filled with lubricating ointment (Paralub Vet Ointment). At the end of each recording session, the chamber was flushed, filled with fresh ointment, and capped with UV-cured cement (Flow-It ALC). At the conclusion of each recording, the chamber was flushed, filled with new ointment, and capped with UV-cured cement.

A 64-channel silicon probe (H3, Cambridge Neurotech) was slowly advanced (100 mm/s) into A1 perpendicular to the pial surface until the tip of the electrode was 1.3-1.4mm below the cortical surface to cover all layers of A1. The brain was allowed to settle for at least 15 min before recordings began. On the day of the first recording, multiple penetrations were made to identify the tonotopic reversal which represents the rostral border of A1 (Guo et al. 2012). Raw neural data was digitized at 32-bit, 24.4 kHz and stored in binary format (PZ5 Neurodigitizer, RZ2 BioAmp Processor, RS4 Data Streamer; Tucker-Davis Technologies). To eliminate artifacts, the common mode signal (channel-averaged neural traces) was subtracted from all channels in the brain. Signals were notch filtered at 60 Hz, band-pass filtered (300-3000 Hz, second order Butterworth filter). To calculate local field potentials (LFP), raw signals were first notch filtered at 60 Hz and downsampled to 1 kHz. The current source density (CSD) was calculated as the second spatial derivative of the LFP signal. To eliminate potential artifacts introduced by impedance mismatching between channels, signals were spatially smoothed along the channels with a triangle filter (5-point Hanning window). Two CSD signatures were used to identify L4 in accordance with prior studies: a brief current sink first occurs approximately 10 ms after the onset of a broadband noise burst, which was used to determine the lower border of L4 (Kaur et al. 2005) (50 ms duration, 70 dB SPL, 50 trials). A tri-phasic CSD pattern (sink-source-sink from upper to lower channels) occurs between 20 ms and 50 ms, where the border between the upper sink and the source was used to define the upper boundary of L4 (Müller-Preuss and Mitzdorf 1984; Metherate et al. 2005; Guo et al. 2017; Clayton et al. 2021). Single unit clusters were obtained using Kilosort (Pachitariu et al. 2016), and single unit isolation was based on the presence of both a refractory period within the interspike interval histogram, and an isolation distance (> 10) indicating that single-unit clusters were separated from the surrounding noise (Schmitzer-Torbert et al. 2005; Clayton et al. 2021). BFCN axon terminals were stimulated via an optic fiber/ferrule assembly (0.2 mm diameter, 0.22 NA Doric) coupled to a 473 nm diode laser for ChR2 activation (Omnicron LuxX) at 2.55 mW/mm^2^.

### *In vitro* electrophysiology

Four weeks after viral delivery of ChR2, mice were anesthetized with 5% isoflurane in O2 followed by intraperitoneal administration of Fatal Plus (0.22 ml/kg). Immediately after anesthesia, mice were perfused with ice cold slicing artificial cerebrospinal fluid (ACSF) containing (in mM): 160 sucrose, 28 NaHCO3, 2.5 KCl, 1.25 NaH2PO4, 7.25 glucose, 20 HEPES, 3 Na-pyruvate, 3 Na-ascorbate, 7.5 MgCl2, 1 CaCl2 (Vattino et al. 2024). After perfusion, mice were decapitated and the brain quickly removed. Thalamocortical slices (300 µm) containing the right A1 were obtained by slicing at an angle of 15° from horizontal view (Vattino et al. 2024) and collected in ice cold slicing ACSF on a vibrating microtome (Leica Microsystems; VT1200S). Slices were placed for 30 minutes at 35° C in a chamber filled with recovery ACSF, containing (in mM): 92 NaCl, 28.5 NaHCO3, 2.5 KCl, 1.2 NaH2PO4, 25 glucose, 20 HEPES, 3 Na-pyruvate, 5 Na-ascorbate, 4 MgCl2, 2 CaCl2 (Vattino et al. 2024). After recovery, slices were transferred to a chamber with recording ACSF, containing (in mM): 125 NaCl, 2.5 KCl, 1.25 NaH2PO4, 25 NaHCO3, 25 glucose, 1 MgCl2, 2 CaCl2 (Vattino et al. 2024). Slices were moved to the recording chamber and maintained under continuous superfusion with recording ACSF at near-physiological temperature (31-33° C) during recordings. All solutions used were continuously bubbled with CO2-O2 (95%-5%).

Recording electrodes (3–5 MΩ) were obtained from borosilicate glass capillaries pulled with a micropipette puller (P-97, Sutter Instrument) and filled with current-clamp internal solution, containing (in mM): 5 KCl, 127.5 K-gluconate, 10 HEPES, 2 MgCl2, 0.6 EGTA, 2 Mg-ATP, 0.3 Na-GTP, 5 Na2-Phosphocreatine; pH 7.2, adjusted with KOH (Takesian et al. 2013). The initial series resistance of all recorded cells was below 30 MΩ and compensated up to 60%. Data were collected at a sampling rate of 10 kHz with a Multiclamp 700B amplifier (Molecular Devices), low-pass filtered at 3 kHz and digitized with a digital-to-analog converter board (National Instruments, NI-USB-6343). Custom-designed MATLAB 2018 software was used for data acquisition. All experiments were carried out in a motorized upright microscope (Scientifica, SliceScope Pro 1000) coupled to a CCD camera (Hamamatsu Photonics, Orca Flash 4.0).

Optogenetic stimulation of BFCN axons in A1 was performed with a wide-field 470 nm LED pulse (CoolLED, pE-100; 5 ms, ∼14 mW/mm^2^). ACh-mediated excitatory or inhibitory postsynaptic potentials (E/IPSPs) were obtained from visually identified L1-INs or L6-PNs under IR-DIC by recording in current clamp configuration, in the presence of 20 μM DNQX and 50 μM AP5 to prevent glutamatergic-mediated responses. The cholinergic nature of these responses was further confirmed by bath application of nAChR antagonists methyllycaconitine (MLA, 10 μM) and Dihydro-β-erythroidine hydrobromide (DhβE, 10 μM), or the mAChR antagonist atropine (10 μM).

### Quantification and statistical analysis

For single unit recordings, analysis was performed using custom routines written in MATLAB 2021 (MathWorks). Units were classified based on the ratio of the mean trough-to-peak interval in regular spiking (RS, > 0.6 ms) or fast spiking (FS, < 0.5 ms). For each unit, the average of 50 trials was z-scored and plotted in neurograms (**Figure 4D**). The average z-scored values in **Figure 4E** for each unit were calculated as follows: for the baseline time period, the time bins of the average z-scored response were shuffled 1000 times, three consecutive bins were randomly picked and averaged to obtain 1000 average values for each unit, which were then averaged to obtain a unique value; for the ‘onset’ time period, the three bins corresponding to this time domain were averaged together; for the ‘persistent’ time period, the bin with the largest z-score value was found and averaged with the two adjacent bins. For **Figure 4F**, the values for ‘L2-5’ and ‘L6’ are those corresponding to ‘onset’ in **Figure 4E**, whereas the shuffled control values were obtained for each unit by shuffling the time bins of the entire neurogram 1000 times and averaging the three bins corresponding to the time of the laser stimulation. For *in vitro* whole-cell recordings, excitatory and inhibitory postsynaptic potential amplitudes were quantified using Clampfit 10.7 (Molecular Devices). Statistical comparisons were performed in GraphPad Prism 10. Data represents mean ± SEM unless otherwise noted.

## REFERENCES

1. Abs E, Poorthuis RB, Apelblat D, Muhammad K, Pardi MB, Enke L, Kushinsky D, Pu D-L, Eizinger MF, Conzelmann K-K, Spiegel I, Letzkus JJ. 2018. Learning-Related Plasticity in Dendrite-Targeting Layer 1 Interneurons. Neuron. 100:1–16.

2. Albuquerque EX, Pereira EFR, Alkondon M, Rogers SW. 2009. Mammalian nicotinic acetylcholine receptors: from structure to function. Physiol Rev. 89:73–120.

3. Allaway KC, Muñoz W, Tremblay R, Sherer M, Herron J, Rudy B, Machold R, Fishell G. 2020. Cellular birthdate predicts laminar and regional cholinergic projection topography in the forebrain. Elife. 9.

4. Ananth MR, Rajebhosale P, Kim R, Talmage DA, Role LW. 2023. Basal forebrain cholinergic signalling: development, connectivity and roles in cognition. Nat Rev Neurosci. 24:233– 251.

5. Arroyo S, Bennett C, Aziz D, Brown SP, Hestrin S. 2012. Prolonged disynaptic inhibition in the cortex mediated by slow, non-α7 nicotinic excitation of a specific subset of cortical interneurons. J Neurosci. 32:3859–3864.

6. Asokan MM, Watanabe Y, Kimchi EY, Polley DB. 2023. Potentiation of cholinergic and corticofugal inputs to the lateral amygdala in threat learning. Cell Rep. 42:113167.

7. Bakin JS, Weinberger NM. 1996. Induction of a physiological memory in the cerebral cortex by stimulation of the nucleus basalis. Proc Natl Acad Sci U S A. 93:11219–11224.

8. Bloem B, Schoppink L, Rotaru DC, Faiz A, Hendriks P, Mansvelder HD, van de Berg WDJ, Wouterlood FG. 2014. Topographic mapping between basal forebrain cholinergic neurons and the medial prefrontal cortex in mice. J Neurosci. 34:16234–16246.

9. Bortone DS, Olsen SR, Scanziani M. 2014. Translaminar inhibitory cells recruited by layer 6 corticothalamic neurons suppress visual cortex. Neuron. 82:474–485.

10. Brombas A, Fletcher LN, Williams SR. 2014. Activity-dependent modulation of layer 1 inhibitory neocortical circuits by acetylcholine. J Neurosci. 34:1932–1941.

11. Carr DB, Surmeier DJ. 2007. M1 muscarinic receptor modulation of Kir2 channels enhances temporal summation of excitatory synaptic potentials in prefrontal cortex pyramidal neurons. J Neurophysiol. 97:3432–3438.

12. Chavez C, Zaborszky L. 2017. Basal forebrain cholinergic-auditory cortical network: Primary versus nonprimary auditory cortical areas. Cereb Cortex. 27:2335–2347.

13. Clayton KK, Williamson RS, Hancock KE, Tasaka G-I, Mizrahi A, Hackett TA, Polley DB. 2021. Auditory Corticothalamic Neurons Are Recruited by Motor Preparatory Inputs. Curr Biol. 31:310–321.e5.

14. Cohen-Kashi Malina K, Tsivourakis E, Kushinsky D, Apelblat D, Shtiglitz S, Zohar E, Sokoletsky M, Tasaka G-I, Mizrahi A, Lampl I, Spiegel I. 2021. NDNF interneurons in layer 1 gain-modulate whole cortical columns according to an animal’s behavioral state. Neuron. 109:2150–2164.e5.

15. Couey JJ, Meredith RM, Spijker S, Poorthuis RB, Smit AB, Brussaard AB, Mansvelder HD. 2007. Distributed network actions by nicotine increase the threshold for spike-timing-dependent plasticity in prefrontal cortex. Neuron. 54:73–87.

16. Crandall SR, Cruikshank SJ, Connors BW. 2015. A corticothalamic switch: controlling the thalamus with dynamic synapses. Neuron. 86:768–782.

17. Crouse RB, Kim K, Batchelor HM, Girardi EM, Kamaletdinova R, Chan J, Rajebhosale P, Pittenger ST, Role LW, Talmage DA, Jing M, Li Y, Gao X-B, Mineur YS, Picciotto MR. 2020. Acetylcholine is released in the basolateral amygdala in response to predictors of reward and enhances the learning of cue-reward contingency. Elife. 9.

18. Disney AA, Higley MJ. 2020. Diverse Spatiotemporal Scales of Cholinergic Signaling in the Neocortex. J Neurosci. 40:712–719.

19. Eggermann E, Kremer Y, Crochet S, Petersen CCH. 2014. Cholinergic signals in mouse barrel cortex during active whisker sensing. Cell Rep. 9:1654–1660.

20. Frandolig JE, Matney CJ, Lee K, Kim J, Chevée M, Kim S-J, Bickert AA, Brown SP. 2019. The synaptic organization of layer 6 circuits reveals inhibition as a major output of a neocortical sublamina. Cell Rep. 28:3131–3143.

21. Froemke RC, Carcea I, Barker AJ, Yuan K, Seybold BA, Martins ARO, Zaika N, Bernstein H, Wachs M, Levis PA, Polley DB, Merzenich MM, Schreiner CE. 2013. Long-term modification of cortical synapses improves sensory perception. Nat Neurosci. 16:79–88.

22. Froemke RC, Merzenich MM, Schreiner CE. 2007. A synaptic memory trace for cortical receptive field plasticity. Nature. 450:425–429.

23. Fu Y, Tucciarone JM, Sebastian Espinosa J, Sheng N, Darcy DP, Nicoll RA, Josh Huang Z, Stryker MP. 2014. A Cortical Circuit for Gain Control by Behavioral State. Cell. 156:1139–1152.

24. Ghamari-Langroudi M, Bourque CW. 2004. Muscarinic receptor modulation of slow afterhyperpolarization and phasic firing in rat supraoptic nucleus neurons. J Neurosci. 24:7718–7726.

25. Ghimire M, Cai R, Ling L, Brownell KA, Wisner KW, Cox BC, Hackett TA, Brozoski TJ, Caspary DM. 2023. Desensitizing nicotinic agents normalize tinnitus-related inhibitory dysfunction in the auditory cortex and ameliorate behavioral evidence of tinnitus. Front Neurosci. 17:1197909.

26. Ghimire M, Cai R, Ling L, Hackett TA, Caspary DM. 2020. Nicotinic receptor subunit distribution in auditory cortex: Impact of aging on receptor number and function. J Neurosci. 40:5724–5739.

27. Giessel AJ, Sabatini BL. 2010. M1 muscarinic receptors boost synaptic potentials and calcium influx in dendritic spines by inhibiting postsynaptic SK channels. Neuron. 68:936–947.

28. Goard M, Dan Y. 2009. Basal forebrain activation enhances cortical coding of natural scenes. Nat Neurosci. 12:1444–1449.

29. Gritton HJ, Howe WM, Mallory CS, Hetrick VL, Berke JD, Sarter M. 2016. Cortical cholinergic signaling controls the detection of cues. Proc Natl Acad Sci U S A. 113:1089–1097.

30. Guo W, Chambers AR, Darrow KN, Hancock KE, Shinn-Cunningham BG, Polley DB. 2012. Robustness of cortical topography across fields, laminae, anesthetic states, and neurophysiological signal types. J Neurosci. 32:9159–9172.

31. Guo W, Clause AR, Barth-Maron A, Polley DB. 2017. A corticothalamic circuit for dynamic switching between feature detection and discrimination. Neuron. 95:180–194.

32. Guo W, Robert B, Polley DB. 2019. The Cholinergic Basal Forebrain Links Auditory Stimuli with Delayed Reinforcement to Support Learning. Neuron. 103:1164–1177.e6.

33. Hangya B, Ranade SP, Lorenc M, Kepecs A. 2015. Central Cholinergic Neurons Are Rapidly Recruited by Reinforcement Feedback. Cell. 162:1155–1168.

34. Harris KD, Shepherd GMG. 2015. The neocortical circuit: themes and variations. Nat Neurosci. 18:170–181.

35. Hartung J, Schroeder A, Vázquez RAP, Poorthuis RB, Letzkus JJ. 2024. Layer 1 NDNF Interneurons are Specialized Top-Down Master Regulators of Cortical Circuits. Cell Rep. 23:114212.

36. Hay YA, Lambolez B, Tricoire L. 2016. Nicotinic transmission onto layer 6 cortical neurons relies on synaptic activation of non-α7 receptors. Cereb Cortex. 26:2549–2562.

37. Higley MJ, Picciotto MR. 2014. Neuromodulation by acetylcholine: examples from schizophrenia and depression. Curr Opin Neurobiol. 29:88–95.

38. Kassam SM, Herman PM, Goodfellow NM, Alves NC, Lambe EK. 2008. Developmental excitation of corticothalamic neurons by nicotinic acetylcholine receptors. J Neurosci. 28:8756–8764.

39. Kaur S, Rose HJ, Lazar R, Liang K, Metherate R. 2005. Spectral integration in primary auditory cortex: laminar processing of afferent input, in vivo and in vitro. Neuroscience. 134:1033–1045.

40. Kilgard MP, Merzenich MM. 1998. Cortical map reorganization enabled by nucleus basalis activity. Science. 279:1714–1718.

41. Kim J, Matney CJ, Blankenship A, Hestrin S, Brown SP. 2014. Layer 6 corticothalamic neurons activate a cortical output layer, layer 5a. J Neurosci. 34:9656–9664.

42. Kim J-H, Jung A-H, Jeong D, Choi I, Kim K, Shin S, Kim SJ, Lee S-H. 2016. Selectivity of neuromodulatory projections from the basal forebrain and locus ceruleus to primary sensory cortices. J Neurosci. 36:5314–5327.

43. Kimchi EY, Burgos-Robles A, Matthews GA, Chakoma T, Patarino M, Weddington JC, Siciliano C, Yang W, Foutch S, Simons R, Fong M-F, Jing M, Li Y, Polley DB, Tye KM. 2024. Reward contingency gates selective cholinergic suppression of amygdala neurons. Elife. 12.

44. Kuchibhotla KV, Gill JV, Lindsay GW, Papadoyannis ES, Field RE, Sten TAH, Miller KD, Froemke RC. 2017. Parallel processing by cortical inhibition enables context-dependent behavior. Nat Neurosci. 20:62–71.

45. Laszlovszky T, Schlingloff D, Hegedüs P, Freund TF, Gulyás A, Kepecs A, Hangya B. 2020. Distinct synchronization, cortical coupling and behavioral function of two basal forebrain cholinergic neuron types. Nat Neurosci. 23:992–1003.

46. Ledderose JMT, Zolnik TA, Toumazou M, Trimbuch T, Rosenmund C, Eickholt BJ, Jaeger D, Larkum ME, Sachdev RNS. 2023. Layer 1 of somatosensory cortex: an important site for input to a tiny cortical compartment. Cereb Cortex. 33:11354–11372.

47. Letzkus JJ, Wolff SBE, Meyer EMM, Tovote P, Courtin J, Herry C, Lüthi A. 2011. A disinhibitory microcircuit for associative fear learning in the auditory cortex. Nature. 480:331–335.

48. McGinley MJ, David SV, McCormick DA. 2015. Cortical membrane potential signature of optimal states for sensory signal detection. Neuron. 87:179–192.

49. Mechawar N, Cozzari C, Descarries L. 2000. Cholinergic innervation in adult rat cerebral cortex: a quantitative immunocytochemical description. J Comp Neurol. 428:305–318.

50. Metherate R, Kaur S, Kawai H, Lazar R, Liang K, Rose HJ. 2005. Spectral integration in auditory cortex: mechanisms and modulation. Hear Res. 206:146–158.

51. Müller-Preuss P, Mitzdorf U. 1984. Functional anatomy of the inferior colliculus and the auditory cortex: current source density analyses of click-evoked potentials. Hearing Research. 16:133142.

52. Nelson A, Mooney R. 2016. The Basal Forebrain and Motor Cortex Provide Convergent yet Distinct Movement-Related Inputs to the Auditory Cortex. Neuron. 90:635–648.

53. Obermayer J, Verhoog MB, Luchicchi A, Mansvelder HD. 2017. Cholinergic Modulation of Cortical Microcircuits Is Layer-Specific: Evidence from Rodent, Monkey and Human Brain. Front Neural Circuits. 11:100.

54. Olsen SR, Bortone DS, Adesnik H, Scanziani M. 2012. Gain control by layer six in cortical circuits of vision. Nature. 483:47–52.

55. Pachitariu M, Steinmetz N, Kadir S, Carandini M, Kenneth D. H. 2016. Kilosort: realtime spike-sorting for extracellular electrophysiology with hundreds of channels. bioRxiv. 10.1101/061481

56. Parikh V, Kozak R, Martinez V, Sarter M. 2007. Prefrontal acetylcholine release controls cue detection on multiple timescales. Neuron. 56:141–154.

57. Pi H-J, Hangya B, Kvitsiani D, Sanders JI, Huang ZJ, Kepecs A. 2013. Cortical interneurons that specialize in disinhibitory control. Nature. 503:521–524.

58. Picciotto MR, Higley MJ, Mineur YS. 2012. Acetylcholine as a neuromodulator: cholinergic signaling shapes nervous system function and behavior. Neuron. 76:116–129.

59. Pinto L, Goard MJ, Estandian D, Xu M, Kwan AC, Lee S-H, Harrison TC, Feng G, Dan Y. 2013. Fast modulation of visual perception by basal forebrain cholinergic neurons. Nat Neurosci. 16:1857–1863.

60. Poorthuis RB, Bloem B, Schak B, Wester J, de Kock CPJ, Mansvelder HD. 2013. Layer-specific modulation of the prefrontal cortex by nicotinic acetylcholine receptors. Cereb Cortex. 23:148–161.

61. Poorthuis RB, Muhammad K, Wang M, Verhoog MB, Junek S, Wrana A, Mansvelder HD, Letzkus JJ. 2018. Rapid Neuromodulation of Layer 1 Interneurons in Human Neocortex. Cell Rep. 23:951–958.

62. Prieto JJ, Winer JA. 1999. Layer VI in cat primary auditory cortex: Golgi study and sublaminar origins of projection neurons. J Comp Neurol. 404:332–358.

63. Qi G, Yang D, Messore F, Bast A, Yáñez F, Oberlaender M, Feldmeyer D. 2025. FOXP2- immunoreactive corticothalamic neurons in neocortical layers 6a and 6b are tightly regulated by neuromodulatory systems. iScience. 28:111646.

64. Radnikow G, Feldmeyer D. 2018. Layer- and cell type-specific modulation of excitatory neuronal activity in the neocortex. Front Neuroanat. 12:1.

65. Reimer J, McGinley MJ, Liu Y, Rodenkirch C, Wang Q, McCormick DA, Tolias AS. 2016. Pupil fluctuations track rapid changes in adrenergic and cholinergic activity in cortex. Nat Commun. 7:13289.

66. Robert B, Kimchi EY, Watanabe Y, Chakoma T, Jing M, Li Y, Polley DB. 2021. A functional topography within the cholinergic basal forebrain for encoding sensory cues and behavioral reinforcement outcomes. Elife. 10.

67. Sabec MH, Wonnacott S, Warburton EC, Bashir ZI. 2018. Nicotinic Acetylcholine Receptors Control Encoding and Retrieval of Associative Recognition Memory through Plasticity in the Medial Prefrontal Cortex. Cell Rep. 22:3409–3415.

68. Sarter M, Lustig C. 2020. Forebrain Cholinergic Signaling: Wired and Phasic, Not Tonic, and Causing Behavior. J Neurosci. 40:720–725.

69. Sarter M, Parikh V, Howe WM. 2009. Phasic acetylcholine release and the volume transmission hypothesis: time to move on. Nat Rev Neurosci. 10:383–390.

70. Schlingloff D, Szabó Í, Gulyás É, Király B, Kispál R, Stephenson-Jones M, Hangya B. 2025. Most ventral pallidal cholinergic neurons are cortically projecting bursting basal forebrain cholinergic neurons. BioRxiv. 10.1101/2025.02.23.639747

71. Schmitzer-Torbert N, Jackson J, Henze D, Harris K, Redish AD. 2005. Quantitative measures of cluster quality for use in extracellular recordings. Neuroscience. 131:1–11.

72. Schofield BR. 2009. Projections to the inferior colliculus from layer VI cells of auditory cortex. Neuroscience. 159:246–258.

73. Sundberg SC, Lindström SH, Sanchez GM, Granseth B. 2018. Cre-expressing neurons in visual cortex of Ntsr1-Cre GN220 mice are corticothalamic and are depolarized by acetylcholine. J Comp Neurol. 526:120–132.

74. Takesian AE, Bogart LJ, Lichtman JW, Hensch TK. 2018. Inhibitory circuit gating of auditory critical-period plasticity. Nat Neurosci. 21:218–227.

75. Takesian AE, Kotak VC, Sharma N, Sanes DH. 2013. Hearing loss differentially affects thalamic drive to two cortical interneuron subtypes. J Neurophysiol. 110:999–1008.

76. Thiele A. 2013. Muscarinic signaling in the brain. Annu Rev Neurosci. 36:271–294.

77. Tremblay R, Lee S, Rudy B. 2016. GABAergic Interneurons in the Neocortex: From Cellular Properties to Circuits. Neuron. 91:260–292.

78. Vattino LG, MacGregor CP, Liu CJ, Sweeney CG, Takesian AE. 2024. Primary auditory thalamus relays directly to cortical layer 1 interneurons. bioRxiv. 10.1101/2024.07.16.603741

79. Verhoog MB, Obermayer J, Kortleven CA, Wilbers R, Wester J, Baayen JC, De Kock CPJ, Meredith RM, Mansvelder HD. 2016. Layer-specific cholinergic control of human and mouse cortical synaptic plasticity. Nat Commun. 7:1–13.

80. Voigts J, Deister CA, Moore CI. 2020. Layer 6 ensembles can selectively regulate the behavioral impact and layer-specific representation of sensory deviants. Elife. 9.

81. West DC, Mercer A, Kirchhecker S, Morris OT, Thomson AM. 2006. Layer 6 cortico-thalamic pyramidal cells preferentially innervate interneurons and generate facilitating EPSPs. Cereb Cortex. 16:200–211.

82. Williams SR, Fletcher LN. 2019. A Dendritic Substrate for the Cholinergic Control of Neocortical Output Neurons. Neuron. 101:486–499.

83. Williamson RS, Polley DB. 2019. Parallel pathways for sound processing and functional connectivity among layer 5 and 6 auditory corticofugal neurons. Elife. 8.

84. Winer JA, Prieto JJ. 2001. Layer V in cat primary auditory cortex (AI): cellular architecture and identification of projection neurons. J Comp Neurol. 434:379–412.

85. Womble MD, Moises HC. 1992. Muscarinic inhibition of M-current and a potassium leak conductance in neurones of the rat basolateral amygdala. J Physiol. 457:93–114.

86. Yaeger CE, Ringach DL, Trachtenberg JT. 2019. Neuromodulatory control of localized dendritic spiking in critical period cortex. Nature. 567:100–104.

87. Yang D, Günter R, Qi G, Radnikow G, Feldmeyer D. 2020. Muscarinic and nicotinic modulation of neocortical layer 6A synaptic microcircuits is cooperative and cell-specific. Cereb Cortex. 30:3528–3542.

88. Yudintsev G, Asilador AR, Sons S, Vaithiyalingam Chandra Sekaran N, Coppinger M, Nair K, Prasad M, Xiao G, Ibrahim BA, Shinagawa Y, Llano DA. 2021. Evidence for layer-specific connectional heterogeneity in the mouse auditory corticocollicular system. J Neurosci. 41:9906–9918.

89. Zhu F, Elnozahy S, Lawlor J, Kuchibhotla KV. 2023. The cholinergic basal forebrain provides a parallel channel for state-dependent sensory signaling to auditory cortex. Nat Neurosci. 26:810–819.

90. Zoli M, Pucci S, Vilella A, Gotti C. 2018. Neuronal and extraneuronal nicotinic acetylcholine receptors. Curr Neuropharmacol. 16:338–349.

