## Supplementary Table 1 for "Layer 6 is a hub for cholinergic modulation in the mouse auditory cortex"

**Supplementary Table 1 – related to Figure 1D**

| **Total number of transcripts (N=3 mice)** | **L1** | **L2/3** | **L4** | **L5** | **L6** |
| --- | --- | --- | --- | --- | --- |
| **α4** | 1715.67  ±  316.53 | 2684.67  ±  331.44 | 3910.67  ±  365.88 | 14232.33  ±  916.99 | 31700.33  ±  1501.88 |
| **α7** | 1456.33  ±  35.04 | 3880.67  ±  696.29 | 5769.67  ±  1362.55 | 6632.00  ±  1337.14 | 12012.00  ±  2770.313 |
| **β2** | 986.67  ±  179.74 | 9845.00  ±  962.41 | 14793.33  ±  2220.30 | 13397.33  ±  1980.20 | 15225.33  ±  1789.46 |
| **M1** | 181.00  ±  46.64 | 21613.67  ±  3688.20 | 18406.33  ±  3303.23 | 13663.33  ±  1201.17 | 22662.67  ±  2629.09 |
| **M2** | 21.67  ±  10.63 | 2267.00  ±  333.16 | 11061.00  ±  906.33 | 5206.67  ±  353.60 | 4875.00  ±  227.50 |
| **M3** | 398.67  ±  64.91 | 13644.33  ±  687.56 | 8208.00  ±  421.91 | 8078.33  ±  514.97 | 16139.00  ±  1185.99 |
| **M4** | 106.67  ±  38.90 | 6569.00  ±  840.07 | 12240.67  ±  2337.579 | 7147.67  ±  1180.10 | 10009.33  ±  1815.24 |
