## Supplementary Table 2 for "Layer 6 is a hub for cholinergic modulation in the mouse auditory cortex"

**Supplementary Table 2 – related to Figure 2B and Figure 3B**

| **Fraction of cells (N = 3 mice)** | | **L1** | **L2/3** | **L4** | **L5** | **L6** |
| --- | --- | --- | --- | --- | --- | --- |
| **VGluT1^+^** | **α4+β2** | - | 2.40 ± 0.36 | 2.99 ± 0.69 | 34.98 ± 1.05 | 26.41 ± 3.17 |
|  | **α7** | - | 0.66 ± 0.23 | 0.34 ± 0.14 | 1.10 ± 0.17 | 1.19 ± 0.10 |
|  | **α4+β2+α7** | - | 0.71 ± 0.58 | 1.91 ± 1.20 | 16.04 ± 4.21 | 29.03 ± 6.68 |
|  | **M1** | - | 87.38 ± 4.06 | 72.83 ± 5.78 | 72.35 ± 5.89 | 78.00 ± 4.74 |
|  | **M2** | - | 10.28 ± 2.48 | 53.79 ± 4.30 | 29.71 ± 3.97 | 13.13 ± 1.27 |
|  | **M3** | - | 69.33 ± 4.79 | 31.87 ± 0.65 | 44.42 ± 1.89 | 62.07 ± 1.64 |
|  | **M4** | - | 32.70 ± 5.51 | 50.46 ± 9.39 | 39.50 ± 5.84 | 36.17 ± 8.50 |
| **VGAT^+^** | **α4+β2** | 12.12 ± 9.90 | 17.10 ± 8.27 | 16.22 ± 1.98 | 17.92 ± 3.57 | 20.02 ± 5.16 |
|  | **α7** | - | - | - | - | - |
|  | **α4+β2+α7** | 75.17 ± 1.19 | 20.41 ± 5.99 | 8.73 ± 3.94 | 11.71 ± 1.54 | 8.91 ± 2.30 |
|  | **M1** | - | 26.20 ± 10.61 | 21.30 ± 9.37 | 14.55 ± 1.52 | 12.45 ± 4.48 |
|  | **M2** | - | 2.22 ± 1.81 | 6.68 ± 2.74 | 14.15 ± 2.03 | 30.28 ± 4.74 |
|  | **M3** | 7.41 ± 6.05 | 31.83 ± 5.56 | 29.58 ± 3.21 | 26.17 ± 3.04 | 37.61 ± 5.69 |
|  | **M4** | - | 14.85 ± 2.95 | 26.22 ± 9.11 | 17.89 ± 3.87 | 29.30 ± 4.53 |
