## Supplementary Table 3 for "Layer 6 is a hub for cholinergic modulation in the mouse auditory cortex"

**Supplementary Table 3 – related to Figure 2C and Figure 3C**

| **Fraction of cells (N = 3 mice)** | **α4β2** | **α7** | **α4+β2+α7** | **M1** | **M2** | **M3** | **M4** |
| --- | --- | --- | --- | --- | --- | --- | --- |
| **CTB^+^** | 29.16  ±  2.99 | - | 33.78  ±  5.98 | 84.75  ±  5.59 | 7.17  ±  1.69 | 68.67  ±  1.70 | 40.86  ±  8.83 |
| **CTB^-^** | 22.21  ±  2.81 | 2.82  ±  0.53 | 22.62  ±  6.38 | 70.64  ±  5.95 | 20.36  ±  3.70 | 55.43  ±  1.91 | 30.86  ±  8.48 |
